# Enhanced sensory coding in mouse vibrissal and visual cortex through TRPA1

**DOI:** 10.1101/2019.12.19.881896

**Authors:** Ehsan Kheradpezhouh, Matthew F. Tang, Jason B. Mattingley, Ehsan Arabzadeh

## Abstract

Transient Receptor Potential Ankyrin 1 (TRPA1) is a non-selective cation channel, which is broadly expressed throughout the body. Despite its expression in the mammalian cortex, little is known about the contribution of TRPA1 to cortical function. Here we investigate the role of TRPA1 in sensory information processing by performing electrophysiological recording and 2-photon calcium imaging from two sensory areas in mice: the primary vibrissal somatosensory cortex (vS1) and the primary visual cortex (V1). In vS1, local activation of TRPA1 by its agonist AITC significantly increased the spontaneous activity of cortical neurons, their evoked response to vibrissal stimulation, and their response range, consistent with a positive gain modulation. TRPA1 inhibition with HC-030031 reversed these modulations to below initial control gains. The gain modulations were absent in TRPA1 Knockout mice. In V1, TRPA1 activation increased the gain of direction and orientation selectivity similarly to the gain modulations observed in vS1 cortex. Linear decoding analysis of V1 population activity confirmed faster and more reliable encoding of visual signals in the presence of TRPA1 activation. Overall, our findings reveal a physiological role for TRPA1 in enhancing sensory signals in the mammalian cortex.

## Introduction

A key challenge in neuroscience is to understand the molecular and circuit mechanisms that underlie efficient sensory processing in the mammalian cortex. A productive approach is to identify the cellular and molecular mechanisms that are preserved across species and to examine how these mechanisms generalize from simpler organisms to more complex brains. The Transient Receptor Potential Ankyrin1 (TRPA1) is a non-selective cation channel that is preserved across species from insects [1] to the mammalian brain [2], with homologues in fungi and algae [3,4]. TRPA1 belongs to the TRP superfamily, an ancient family of channels [5] with diverse roles in sensation and behavior [6]. In simple organisms, such as invertebrates, TRPA1 acts as a polymodal sensor [7] and is directly involved in processing of sensory information across multiple modalities including in detection of noxious chemicals, light and temperature changes [8,9]. In mammals, TRPA1 is found in many organelles and tissues including those of the central nervous system [2]. Our knowledge of the role of TRPA1 in the nervous system has been limited, mostly to its involvement in the pain pathway [10] and, in particular, in dorsal root ganglia (DRG) neurons [10–12]. However, there has been growing evidence indicating a potential role for TRPA1 in the mammalian brain; TRPA1 activation was found to enhance neuronal excitability by increasing intracellular calcium in astrocytes in the hippocampus [13] and motor cortex [14].

Recently, we identified that in both rats and mice, TRPA1 is widely expressed in the primary sensory cortex [15]. Furthermore, we found that cortical neurons were depolarized by *in vitro* activation of TRPA1 [15,16]. However, the role of this channel *in vivo* and its potential contribution to sensory processing in the mammalian cortex remains unknown. Here, we studied TRPA1 modulations of sensory processing in two well-characterized cortical areas in mice: the primary vibrissal (whisker) somatosensory cortex (vS1) and the primary visual cortex (V1). The rodent vS1 provides an excellent model to investigate neuronal coding due to its functional efficiency [17] and structural organization [18–20]. Similarly, the tuning properties of V1 neurons are well-studied and can be characterized in response to controlled stimuli such as drifting gratings [21]. We quantified how TRPA1 activation and suppression modulate baseline neuronal activity and the evoked responses to sensory stimuli using electrophyiological recording and two-photon calcium imaging in vS1 and V1 cortex.

## Results

### TRPA1 modulation of neuronal activity in vS1 cortex

To identify any potential effect of TRPA1 on sensory processing, we employed loose cell-attached recording of individual neurons across layers of the vS1 cortex under local activation or suppression of TRPA1 (Fig. 1A). To limit the pharmacological modulation of TRPA1 to the local area around the recorded neuron, we custom-built a pipette pair with a tip-to-tip distance of ~50 µm (magnified inset, Fig. 1A). We recorded/labelled individual neurons using the high impedance “recording” pipette (narrow tip with 5-10 MΩ resistance) under continual application of aCSF (control), TRPA1 agonist (AITC, 500 µM) or TRPA1 antagonist (HC-030031, 10 µM) through the “infusion” pipette (wide tip, 25-50 µm). We applied a series of deflections to the whisker pad (6 amplitudes from 0 to 200 μm) and studied neuronal responses in the contralateral vS1 under these three conditions. Figure 1B illustrates how TRPA1 modulation affects the spiking activity of an example vS1 neuron in response to a 200-μm deflection. TRPA1 activation increased the neuron’s ongoing activity (before stimulus onset) and its stimulus-evoked response. After subsequent application of TRPA1 blocker, these modulations gradually returned to their initial levels (HC, Fig. 1B). Figure 1C illustrates how TRPA1 modulations affected the neuronal response profile to the whole stimulus set for two example vS1 neurons, along with their reconstructed morphology. In both cases, activation of TRPA1 resulted in an upward shift in neuronal response function, which then returned to its baseline profile or lower after subsequent application of TRPA1 blocker.

**Figure 1.**
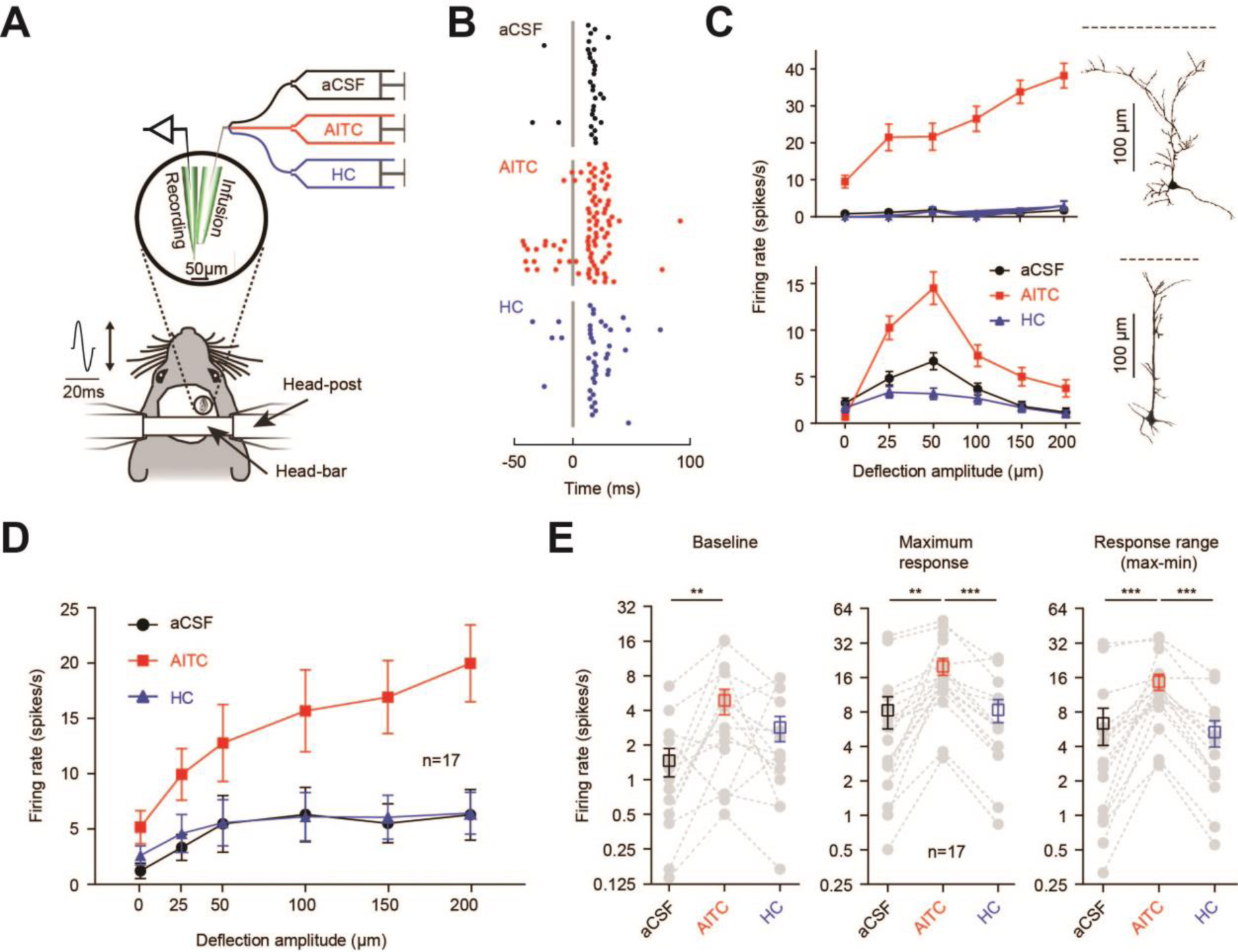
TRPA1 activation increased neuronal response to whisker stimulation. **A.** Schematic custom-made setup for pharmacological manipulation of TRPA1 *in vivo*. The magnified circle shows the pipette pair visualized under a microscope. Single neurons were recorded from vS1 cortex while applying deflections to the contralateral whisker pad. Pharmacological manipulations were applied through the ‘infusion’ pipette. **B.** Raster plots of spiking activity for an example neuron in the presence of aCSF (control), AITC (TRPA1 agonist, 500 µM) and HC (TRPA1 antagonist, 10 µM). **C.** Two sample neurons recorded in layer 2/3 vS1 cortex show modulations in neuronal tuning with TRPA1 activation (AITC) or suppression (HC). The insets show histological reconstruction of each neuron. The horizontal dashed lines indicate cortical surface. **D.** The average neuronal response function across 17 recorded neurons. **E.** Neuronal response is plotted in logarithmic scale. Gray dots represent individual neurons. The black squares represent the average values across the population for baseline (left), peak (middle) and response range (right). Error bars are standard error of the mean across neurons. In all panels, ** *p* < 0.01 and *** *p* < 0.001.

The TRPA1 enhancement of neuronal activity was observed across all recorded neurons (Fig. 1D; n=17). Figure 1E quantifies how TRPA1 activation or suppression affected three key parameters of the neuronal response function: baseline activity, maximum response, and response range (the difference between the maximum and minimum response). Activation of TRPA1 produced a significant increase in baseline activity (amplitude = 0μm; n=17, Fig. 1E, left panel; *p* < 0.01; paired *t* test), a significant increase in the maximum evoked response (*p* < 0.01; Fig. 1E, middle panel), and a significant increase in response range (*p* < 0.001; Fig. 1E, right panel). After subsequent application of TRPA1 inhibitor (HC), all three measures returned to their initial values (HC, Fig. 1E) with no statistically significant differences between HC and control conditions (NS, *p* > 0.05). These results indicate that TRPA1 activation enhanced neuronal activity in vS1 cortex consistent with a multiplicative gain modulation (see below).

In earlier results, we demonstrated that application of TRPA1 agonist enhanced the neuronal response to whisker stimulation. To confirm that the observed enhancement was indeed specifically mediated through TRPA1 gating, we repeated the experiments on TRPA1 knockout (KO) mice that specifically lack this channel. After prolonged application (20 min) of TRPA1 agonist, we did not observe any significant changes in the baseline activity, maximum response, or response range of any recorded vS1 neurons (7 neurons recorded from 3 TRPA1 KO mice; *p* > 0.05; Fig. 2A and 2B).

### Blocking TRPA1 reduces neuronal activity in vS1 cortex

After quantifying TRPA1 modulations, we noticed that for a number of neurons application of HC reduced the maximum evoked response and the response range to values lower than that observed under the control condition (aCSF, see the example neuron in Fig 1D, lower panel). This observation suggests that TRPA1 may have a baseline level of activation during normal physiological conditions. To examine this possibility, we applied the TRPA1 antagonist (HC, 10 µM) in the absence of any initial application of AITC. Introducing this highly-selective TRPA1 blocker to the recorded neurons (n=13) significantly reduced the baseline activity (*p* < 0.05), the maximum evoked response (*p* < 0.01) and the response range (*p* < 0.01; Fig. 2C & 2D). These results further support that TRPA1 has a baseline level of activation and may thus contribute to sensory processing under normal physiological conditions.

**Figure 2.**
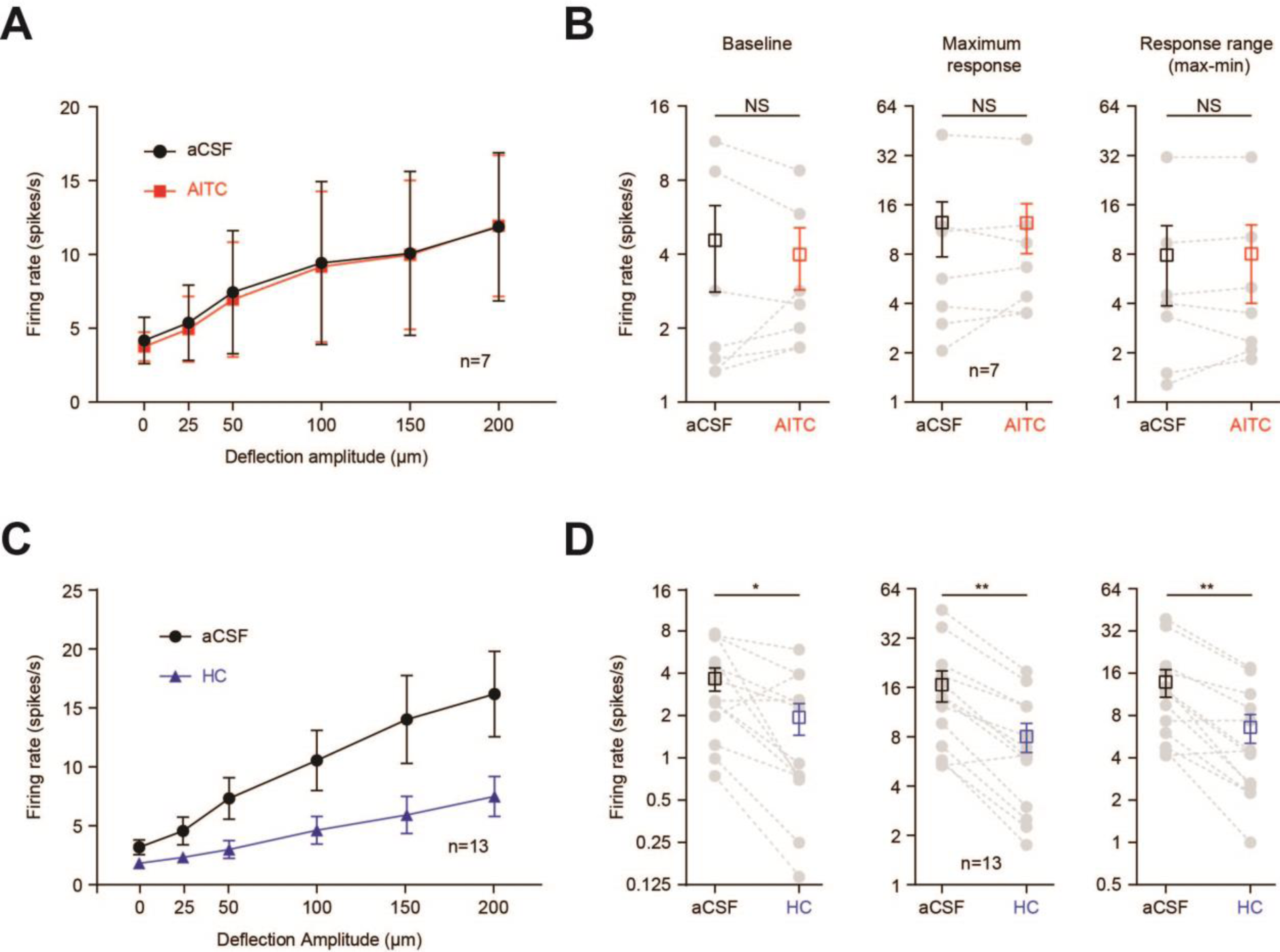
Specificity of the observed modulations to TRPA1. **A.** The average response function of 7 vS1 neurons recorded from 3 TRPA1 KO mice. **B.** In TRPA1 KO mice, AITC produced no statistically significant changes in baseline activity, the peak response or response range of the recorded neurons, *p* > 0.05. **C.** In the absence of any prior activation of TRPA1, application of TRPA1 antagonist (HC, 10 µM) significantly reduced the neuronal response to whisker stimulation. **D.** Application of HC significantly reduced the neuronal baseline activity, the maximum response and the response range. In all panels, error bars are standard error of the mean across neurons. In all panels, * *p* < 0.05, ** *p* < 0.01 and NS not statistically significant.

### TRPA1 activation/suppression produce a gain modulation in neuronal response function

To better understand the effect of TRPA1 modulation on sensory coding, we further quantified the parameters of neuronal response functions. First, we visualized the neuronal response to the full set of stimuli in the presence and absence of TRPA1 modulation. Figure 3A illustrates this form of visualization for the example neuron from Fig. 1C (lower panel). Here, for every stimulus amplitude, the evoked response under aCSF (control) is plotted against the evoked response under AITC (TRPA1 activation). The slope and intercept of the best-fitted line to these data provide key information about the effect of TRPA1 on coding efficiency. The slope captures any multiplicative gain modulations in the neuronal response function, whereas the intercept captures any additive changes in the neuronal response function. For example, a slope of one and a positive intercept would indicate a purely additive modulation, corresponding to a fixed increase in neuronal activity both at baseline and for the evoked responses. On the other hand, a positive slope indicates a multiplicative gain modulation. We applied this quantification to all neurons recorded in wild type and TRPA1 KO mice (Fig. 3). In the wild type mice, TRPA1 activation produced a systematic positive gain modulation in the neuronal response function indicating an enhanced coding capacity (*p* < 0.001, red lines in Fig. 3B) with all recorded neurons exhibiting a slope of >1 (red dots in Fig. 3C). By contrast, TRPA1 inhibition (HC) significantly reduced the gain of the recorded neurons with slopes that were systematically below 1 (p < 0.001, negative gain modulation, blue lines and dots, Figs. 3B and 3C). The intercepts did not show a systematic change with TRPA1 activation or suppression (1.72±1.93 and 0.21±0.54, respectively; NS, *p* > 0.05). Finally, neurons recorded from the TRPA1 KO mice exhibited no systematic changes in slope or intercept (1.02±0.06 and - 0.04±0.4, respectively; red dashed lines and red hollow dots, Figs. 3B and 3C).

**Figure 3.**
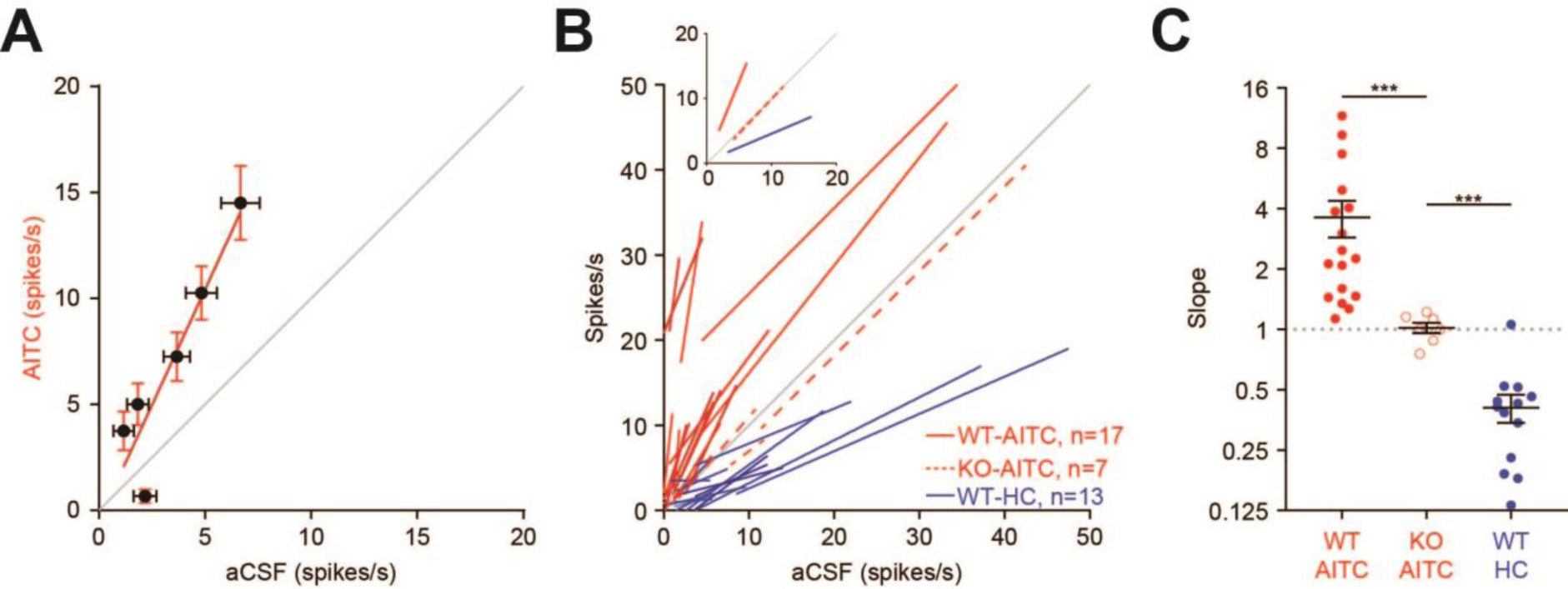
TRPA1 produces a multiplicative gain modulation in vS1 neurons. **A.** Each dot represents the neuronal response of the example neuron (from 1D, lower panel) to a single stimulus amplitude under TRPA1 activation (y-axis, AITC) against control (x-axis, aCSF). The red line represents the best-fitted line (slope = 2.26; intercept = 0.06). Error bars represent standard error of the mean across trials. **B.** Lines are best fits for each recorded neuron. Solid red lines (n=17) represent neurons recorded from wild-type (WT) mice (y-axis, AITC). Dashed red lines (n=7) represent neurons recorded from the TRPA1 KO mice (y-axis, AITC). Blue lines (n=13) represent neurons recorded from WT mice under TRPA1 suppression (y-axis, HC). Note that for all of these neurons the modulation with HC occurred in the absence of any initial activation of TRPA1. The inset shows the best fitted lines for the average response in each of the three populations. **C.** The slopes of the lines presented in **B**. Error bars are standard error of the mean across neurons. *** *p* < 0.001.

### Modulation of cortical population activity by TRPA1

We next examined the effect of TRPA1 modulation on cortical population activity by 2-photon Ca^2+^ imaging of Layer 2/3 neurons. A unique feature of TRPA1 is its high permeability to Ca^2+^; unlike other TRP channels, TRPA1 is more permeable to Ca^2+^ than any other cations (~5.1 times more than Na^+^) [22]. We expressed GCaMP6f (a highly-sensitive Ca^2+^ sensor [23]) in vS1 neurons. After 2-3 weeks, under 2-photon imaging guidance, we inserted an infusion glass pipette close to the target area (Fig. 4B). Similar to earlier experiments, we captured the effect of TRPA1 modulation on the ongoing activity of vS1 neurons and sensory-evoked responses under the control condition (aCSF), TRPA1 activation (AITC, 500 µM) and TRPA1 inhibition (HC, 10 µM). Figure 4C illustrates changes in fluorescence (*ΔF/F0*) in a sample neuron when the whiskers were stimulated at 250 µm intensity. TRPA1 activation increased neuronal fluorescence in response to this whisker stimulation. Subsequent inhibition of TRPA1 with HC reduced *ΔF/F0* to its initial values (Fig. 4C). These findings generalized across the population of neurons imaged in this mouse (n = 87; Fig. 4B). Overall, we imaged the activity of 230 neurons in 4 mice. To better quantify the effect of TRPA1 modulation on neuronal tunings, we plotted the changes in fluorescence over a period of 1 second (area under curve, *ΔF/F0*/s) after stimulation. TRPA1 activation significantly increased neuronal responses to whisker stimulation (n= 230, *p* < 0.01 at 100, 150 and 250 µm stimulation intensities; Fig. 4E). TRPA1 modulations of the evoked response were significantly more pronounced at 250 μm intensity (n = 230, *p* < 0.0001; Fig. 4F). Moreover, TRPA1 activation significantly increased the average fluorescent activity of vS1 neurons during epochs of zero stimulation, suggesting a rise in neuronal Ca^2+^ at the baseline level (t test, *p* value < 0.0001; Fig. 4G). TRPA1 suppression, on the other hand, produced a significant reduction in baseline fluorescence levels (t test, *p* < 0.05; Fig. 4G). Consistent with the earlier electrophysiological findings, the imaging data indicate that TRPA1 activation in vS1 cortex increases neuronal excitability and evoked responses to vibrotactile inputs. Next, we investigated whether the findings we observed in the somatosensory cortex generalize to another well-characterised sensory area, the visual cortex.

**Figure 4.**
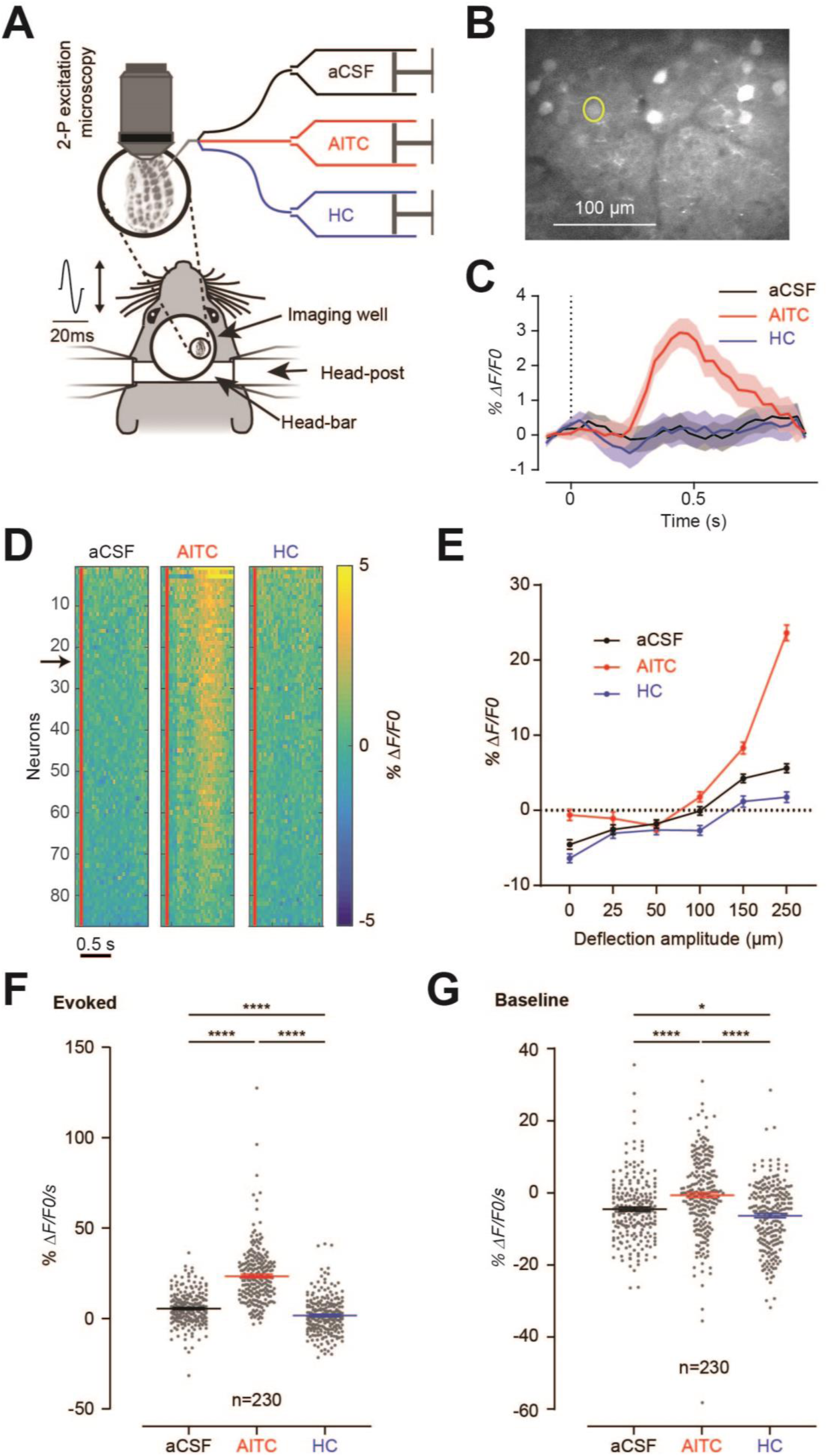
TRPA1 activation enhances neuronal population activity in vS1 cortex. **A.** Schematic setup for 2-photon calcium imaging from mouse vS1 cortex. Similar to earlier experiments, TRPA1 activity was modulated by application of AITC or HC through the infusion pipette. **B.** A 2-photon image of vS1 neurons expressing GCaMP6f. Scale bar is 100 µm. **C.** Changes of fluorescence (*ΔF/F0)* for the neuron marked in **B** in response to 250 µm whisker deflection under control (aCSF), TRPA1 activation (AITC) and TRPA1 inhibition (HC). The dotted line indicates the onset of whisker stimulation. Shaded error bars indicate standard error of mean *ΔF/F0* across trials. **D.** Heat map of neuronal activity across cells shown in **B** (n=87). Every horizontal line represents *ΔF/F0* of a single neuron. Vertical red lines indicate the onset of a 250 µm whisker stimulation. The arrow indicates the example neuron in **C. E.** Changes in *ΔF/F0* are measured over 1-second duration after stimulation onset. **F.** Evoked neuronal response. Every dot represents a neuron. Black lines and error bars indicate the mean and standard error of mean across neurons. Here data are pooled across 4 mice (n = 230 neurons). **G.** Baseline neuronal activity. The change in fluorescence intensity relative to the average whole-session fluorescence. In each condition, the fluorescence is measured for stimulus 0 (no whisker stimulation). In all panels, *, *p* < 0.05; ****, *p* < 0.0001.

### TRPA1 modulates neuronal activity in the mouse visual cortex

We expressed GCaMP6f in L2/3 of the primary visual cortex (V1) and characterized how neuronal tuning to motion direction changed with TRPA1 activation and suppression. For 211 neurons imaged across 3 mice (Fig. 5A), we measured changes in fluorescence (*ΔF/F0*) during repeated presentation of drifting gratings moving in one of 16 different directions (Fig. 5B). Similar to earlier experiments, we quantified the neuronal tuning functions under three conditions: control (aCSF), TRPA1 activation (AITC, 500 µM) and TRPA1 inhibition (HC, 10 µM). Fig. 5C shows how the tuning of a sample neuron was affected by TRPA1 modulation. The neuron showed strong direction tuning with the highest response produced when the stimulus drifted at 292.5° (preferred direction). This neuron was predominantly motion selective as the excitation was approximately half the preferred response when the same orientation drifted in the opposite direction (112.5 degrees). Unsurprisingly for a V1 neuron, the orthogonal orientation (directions of 22.5 and 202.5°) produced the lowest level of activity (polar plot insets, Fig. 5C). TRPA1 activation enhanced the tuning of this neuron without changing its preferred direction (AITC, Fig. 5C). Subsequent application of TRPA1 inhibitor (HC) eliminated the neuronal tuning almost entirely (HC, Fig. 5C). Consistent with previous research [23,24], we identified many neurons with strong direction selectivity across 3 experiments, (n=51; permutation testing, *p* < 0.0001). Figure 5D illustrates the response of each neuron (*ΔF/F0*) to the preferred direction, which was defined as the direction that produced the maximum average response in the 1-second window post stimulus onset. Across the population of imaged neurons, TRPA1 activation significantly increased the response to the preferred direction (n = 51; *p* = 0.0013, TRPA1 vs aCSF).

To better quantify the effect of TRPA1 modulation on neuronal selectivity, we fitted double Gaussian functions [25] to the mean evoked response (0 to 1-second post-stimulus; Fig. 5E). To visualize the population tuning properties, we aligned the tuning curves of all neurons to the same preferred direction (referred to as 0º, Fig. 5E). TRPA1 activation enhanced the average neuronal tuning function. This enhancement was statistically significant at the preferred direction (*p* = 0.006, Figs. 5D, 5E and left panel in 5F) but did not reach statistical significance at the anti-preferred direction (AITC, *p* = 0.145). Subsequent application of TRPA1 inhibitor (HC) reduced the neuronal tuning both to the preferred direction (HC versus AITC *p* < 0.0001; HC versus aCSF, *p* = 0.33, Figs. 5D, 5E and left panel in 5F) and to the anti-preferred direction (HC versus AITC, *p* = 0.011, +180°, Fig. 5E and middle panel in 5F). We did not observe any statistically significant differences in tuning widths among the three conditions (*p* > 0.05 for all, Fig. 5F right panel), although there was a trend for AITC to reduce the mean width, and for HC to increase it (HC – AITC; *p* = .07). Together, these findings suggest that for individual cells and for neuronal populations, TRPA1 activation enhances coding efficiency for visual motion stimuli, whereas TRPA1 inhibition reduces the coding efficiency.

**Figure 5.**
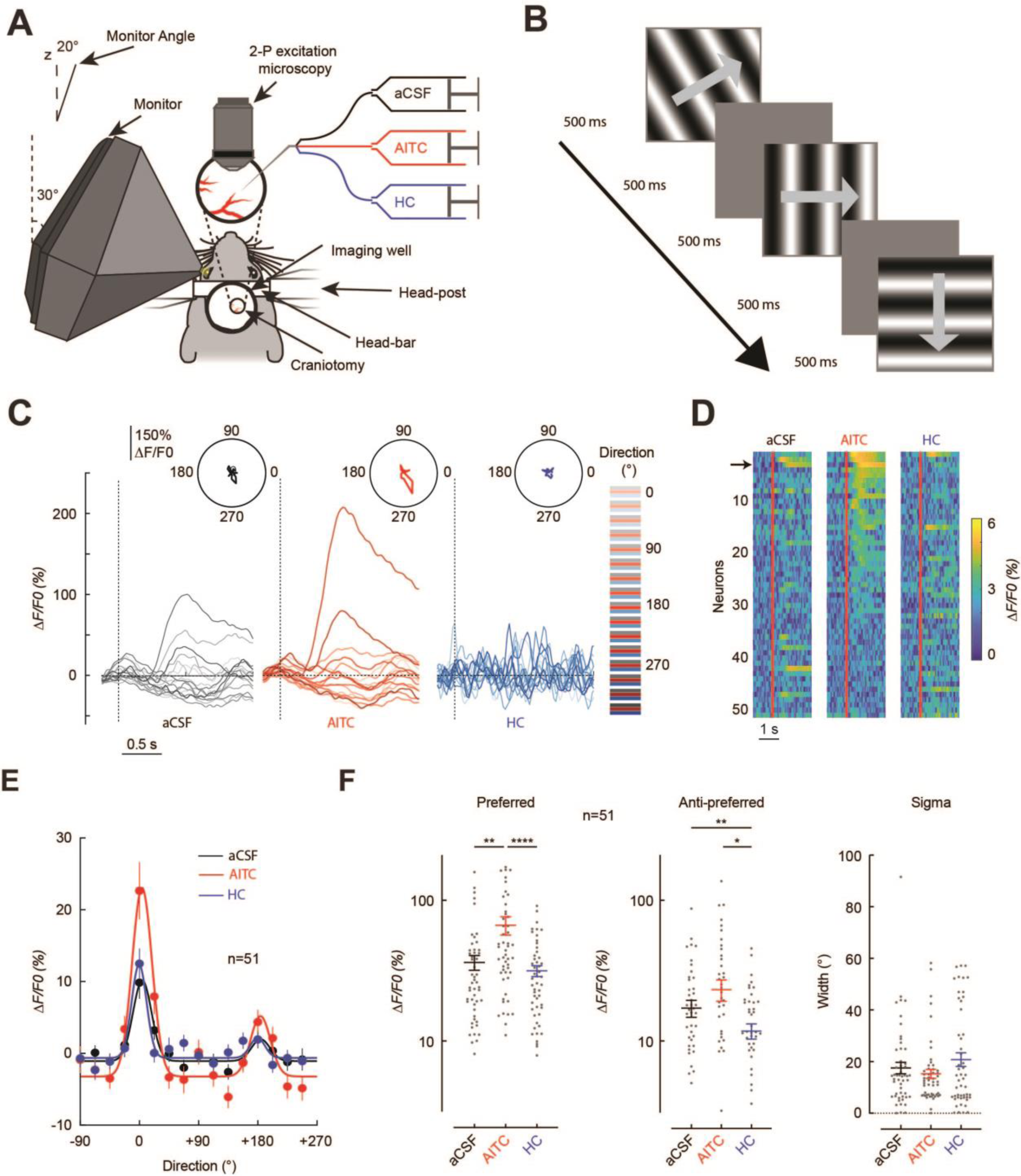
TRPA1 activation increases neuronal response to visual stimulation. **A.** Schematic of the setup for 2-photon calcium imaging from mouse V1 cortex. Similar to earlier experiments, TRPA1 activity was modulated by applying AITC or HC through the infusion pipette. As before, neuronal response was quantified by measuring changes in fluorescent activity of V1 neurons expressing GCaMP6f. To restrict visual stimulation to the contralateral (left) eye, the monitor was mounted on a funnel placed 15 cm from the eye, rotated 30° relative to the dorsoventral axis and tilted 20° vertically. **B.** Drifting gratings were presented in 16 different directions (0° to 337.5° in 22.5° steps) in a pseudorandom sequence and were repeated 35 times for each direction. **C.** Response of a sample neuron to different motion directions. Changes in fluorescence (*ΔF/F0)* were measured relative to the 1-second window prior to stimulus onset (*F0*). Dotted vertical line indicates the stimulus onset. Each line represents the neuronal response to a given stimulus direction. **Insets** show directional tuning in polar plots. **D.** Heat map of neuronal activity in response to the preferred direction. Each horizontal line represents *ΔF/F0* of a single neuron. The arrow indicates the example neuron from **C**. Red lines indicate stimulus onset. **E.** Each dot is the average population response for a stimulus direction (n=51, mean ± SEM). The preferred direction for each neuron was aligned at 0°. Lines indicate double Gaussian fits to mean responses under each condition. **F.** The parameters of the double Gaussian fit for each neuron under the three conditions. Each gray dot represents a neuron. Horizontal lines and error bars indicate the mean and standard error of the mean across neurons. *, *p* < 0.05; **, *p* < 0.01 and ****, *p* < 0.0001.

We next determined how the stimulus representation was affected by TRPA1 at the population level using a multivariate linear discriminant analysis (LDA) approach. The LDA allowed us to quantify the accuracy with which an ideal observer could decode the population response to identify the presented stimulus on a trial-by-trial basis. Figure 6A shows a simplified example of how a simultaneously-imaged neuron pair allowed discrimination between two motion directions. Here, each dot represents the neuronal response in a single trial (35 trials per stimulus). The line that best separates the two stimuli is identified on a subset of ‘training’ trials (black line, Fig. 6A) and is subsequently used to classify stimuli on the remaining ‘test’ trials. This method can be generalized to any population size, and to include all stimulus directions and orientations allowing us to determine how the neuronal population collectively represented the stimulus.

Figure 6B shows decoding accuracy for populations of 15 simultaneously-imaged neurons (n=100 permutations for each of the 3 imaging sessions), under the three experimental conditions (aCSF, AITC, HC). All imaged neurons were included in the permutation regardless of whether they showed direction selectivity. TRPA1 activation significantly improved the population decoding accuracy compared with the control condition (cluster *p* < 0.05, Fig. 6B) whereas TRPA1 inhibition significantly reduced decoding accuracy compared with the control condition (*p* < 0.05, Fig. 6B). In addition to more reliable decoding under TRPA1 activation, the decoding also emerged earlier (87 ms for TRPA1 activation, compared with 203 ms for TRPA1 inhibition).

To determine how these results scale across larger populations of neurons, we applied the same multivariate approach but pooled neurons imaged across three mice (Fig. 6C). The classifier was trained on the average response from 500 to 1000 ms after stimulus presentation (where decoding accuracy was maximal in the previous analysis). This showed that decoding accuracy increased monotonically with the number of neurons (Fig. 6C), with a prominent boost in accuracy with TRPA1 activation (*p* < 0.05). Under TRPA1 inhibition the accuracy was significantly worse relative to control (*p* < 0.05). The modulatory effects of TRPA1 on decoding accuracy became greater with increasing number of neurons. For TRPA1 inhibition, unlike in control and TRPA1 activation, decoding accuracy appears to saturate earlier suggesting that the addition of more neurons did not greatly increase population coding efficiency.

Finally, we investigated how TRPA1 affected the selectivity of cortical neurons for orientation compared with motion direction. Figure 6D summarizes the decoding accuracies for the whole population of imaged neurons. We found that direction decoding (16 directions) led to greater enhancement and degradation with TRPA1 modulation compared with orientation decoding (8 orientations). This effect was confirmed statistically by the presence of a significant interaction in a repeated-measures ANOVA with factors of TRPA1 modulation (control, AITC, HC) and decoding type (motion, orientation), (F(2,27) = 15.12, *p* < 0.001). For stimulus direction, TRPA1 activation and suppression changed the decoding accuracy by +5% and −8.2%, respectively. For stimulus orientation, TRPA1 activation and suppression changed the decoding accuracy by +3.8% and −7.7%, respectively. TRPA1 thus affected direction selectivity more than it affected orientation selectivity (Fig. 6D).

Taken together, as with the results for vS1, we found that TRPA1 activation produced a positive gain modulation in V1 neurons and enhanced their coding efficiency for visual stimuli across the population. The HC-induced reduction in coding efficiency below the control (aCSF) indicates a baseline level of TRPA1 activation under physiological conditions. Collectively, the results from vS1 and V1 suggest a generalized modulatory role for TRPA1 in the processing of sensory signals in the mammalian cortex.

**Figure 6.**
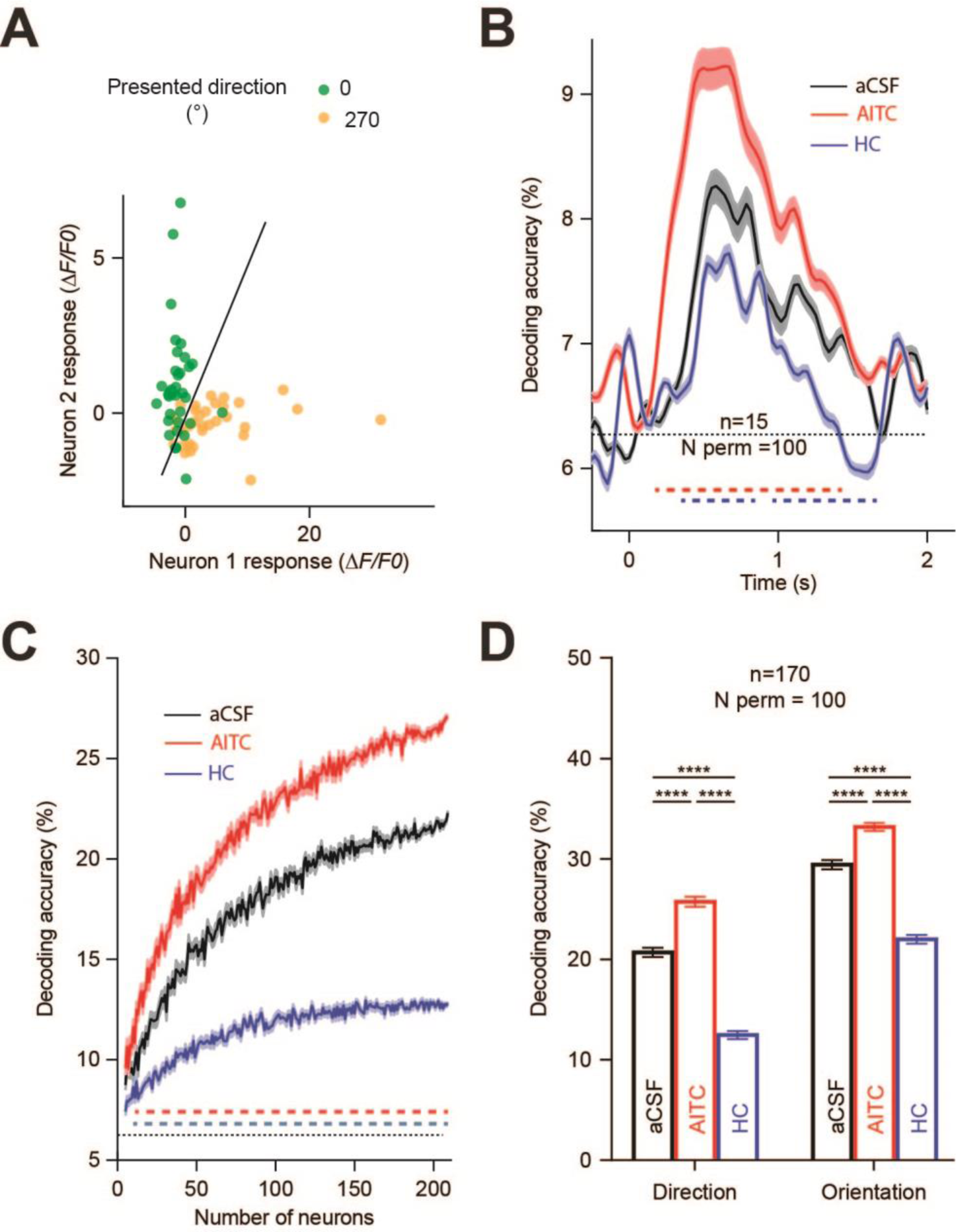
TRPA1 activation improves encoding performance of V1 neurons to visual stimulation. **A.** Example of linear discriminant analysis. Each dot represents one trial. The black line represents the line that best discriminates between the two stimulus responses. **B.** For each of the 3 mice, we randomly selected 100 permutations (N perm) of 15 simultaneously-imaged cells. Decoding accuracy is based on the activity of the 15-neuron population and averaged across permutations. The dotted horizontal line indicates chance level. Decoding accuracy is plotted as a function of time since stimulus onset (0). Horizontal dashed lines show the times at which the difference in decoding accuracy between conditions were statistically significant (comparison of AITC and aCSF is plotted in red; comparison of HC and aCSF is plotted in blue). Accuracy was temporally smoothed with a 43ms kernel Gaussian for display. **C.** Decoding performance for different population sizes. Horizontal dashed lines indicate statistical significance, with conventions as per panel **B**. **D.** Decoding accuracy measured separately for direction (left) and orientation (right) with the classifier trained on mean evoked response (500 to 1000ms). ****, *p* < 0.0001.

## Discussion

TRPA1 belongs to the superfamily of TRP channels, a group of non-selective cation channels that are widely preserved throughout evolution. In simpler organisms, such as invertebrates, TRP channels play a role in sensation of external stimuli including light, sound, chemicals, temperature, and touch [26–30] and are involved in generation of behavioral responses [27,31–34]. Our findings indicate that, as in simpler nervous systems, TRPA1 may contribute to processing of sensory information in the rodent cortex. We characterized how activation and suppression of TRPA1 modulate neuronal responses in mouse somatosensory and visual cortex. In both areas, TRPA1 activation significantly increased the baseline activity of neurons, and produced a positive gain modulation in their evoked response to sensory stimulation. After application of TRPA1 blocker, the neuronal tuning functions returned to their original control profile, or lower, and modulation of tuning was absent in TRPA1 KO mice. We also found that the selective TRPA1 blocker significantly reduced the evoked neuronal responses (negative gain modulation) even in the absence of any initial activation of TRPA1 by the agonist. Together, these findings suggest that cortical TRPA1 has a baseline level of activation under normal physiological conditions.

Understanding the physiological role of TRPA1 in cortical function is challenging, due to the complexity of intracellular regulation of TRPA1 and the fact that endogenous sources of TRPA1 activation and the mechanisms of its desensitization remain uncharacterized [35,36]. TRPA1 is activated by a wide range of exogenous agents. Exogenous pungent molecules such as AITC [37], cinnamaldehyde [38] and allicin [39], and environmental irritants like acroline [40] and 4-Hydroxynonenal (4-HNE) [41] directly activate TRPA1 via interactions with ankyrin repeats (ARs) at the amino-terminal end [42]. Conversely, endogenous compounds including bradykinin [43], histamine [44] and substance-P [39] indirectly activate TRPA1 mainly through G-protein coupled receptors (GPCRs) and phospholipase C (PLC) [43,45]. At present, fundamental questions regarding endogenous activation of TRPA1 remain unaddressed. Recent studies have proposed potential pathways for endogenous TRPA1 activation in the peripheral nervous system and in invertebrates [39,45–47]. TRPA1 is activated by direct interaction with intracellular Ca^2+^ ([Ca^2+^]_i_). As a potent activator, Ca^2+^ was found to directly bind to the C-terminus of TRPA1 opening the pore ([48], but see also [49] for evidence for indirect interaction through calmodulin).

The interaction between GPCRs, PLC and TRPA1 has been studied in the peripheral nervous system. GPCRs, such as the receptors for bradykinin and protease activator receptor 2 (PAR2), can activate TRPA1 and increase its sensitization to other TRPA1 agonists [43,45]. GPCRs also activate PLC which in turn converts phosphatidylinositol 4,5-bisphosphate (PIP2) into inositol trisphosphate (IP3) [50]; PIP2 is a potent TRPA1 blocker and its removal induces further TRPA1 activation [45,50]. In addition, IP3 increases free [Ca^2+^]_i_ which as discussed earlier is a potent TRPA1 activator [50]. TRPA1 is also activated by GPCRs through the second-messenger signalling cascades, including protein kinase A (PKA) and diacylglycerol (DAG) [12,39,45]. In mammals, more than 90% of GPCRs are expressed in the brain [51]. In cortical neurons, GPCRs are expressed in both presynaptic and postsynaptic membrane terminals as well as in astrocytes, where they modulate neurotransmitter release [51,52]. We know that endogenous activation of TRPA1 can result in 30-40% elevation in intracellular calcium [53]. This degree of change in [Ca^2+^]_i_ is expected to dramatically influence neuronal excitability and modulate detection and discrimination of sensory stimuli [54]. Given the high level of GPCR expression in the mammalian brain, including in cortical neurons [52], TRPA1 activation through GPCR cascades provides a potential endogenous mechanism for modulation of sensory processing and behavior under physiological conditions.

The gain modulations induced by TRPA1 are reminiscent of changes induced in neuronal response function induced by attention and neuromodulation. Two key sources of modulation in sensory cortex are the cholinergic system originating from basal forebrain (BF) and noradrenergic system originating from locus ceruleus (LC) [55,56]. Cholinergic and noradrenergic fibres widely project to cortical layers and underlie modulations induced by arousal, and attention [55,56]. Although the mechanisms of these modulations are not fully understood, it is well-known that cholinergic and noradrenergic effects are mediated through GPCRs [55,56]. Muscarinic 1 receptor (M1R) is the most abundant cholinergic receptor in the mammalian cortex [57,58]; during activation by acetylcholine, M1R transforms Gq/11 GPCR which in turn activates PLC enhancing sensory processing [57,59,60]. The noradrenergic system induces its neuromodulation through α1 receptor by transforming Gq GPCR which activates PLC [61,62], and through β receptor by transforming Gs GPCR which activates PKA [63,64]. Given that PLC and PKA are two potential endogenous activators of TRPA1, it is possible that cholinergic and noradrenergic systems mediate their neuromodulation effects partially through cortical TRPA1.

The cellular mechanisms through which TRPA1 modulates evoked responses in sensory cortex remain elusive [65]. Our current results cannot determine the relative contribution of neuronal and astrocytic TRPA1 to the observed modulations in the rodent cortex. There is growing evidence for TRPA1 modulation of neuronal activity in the mammalian brain. In the hippocampus and motor cortex, TRPA1 was found to increase neuronal responses through enhancing astrocytic calcium [13,14]. Oh and colleagues used low-intensity low-frequency ultrasound to directly activate TRPA1 in the motor cortex. Activation of astrocytic TRPA1 in neuron-astrocyte cocultures produced neuronal excitation through increasing astrocytic calcium [13]. Similarly, in rat hippocampal astrocyte-neuron cocultures, TRPA1 activation produced spotty calcium signals in astrocytes which in turn increased neuronal excitability by modulating the efficacy of the interneuron inhibitory synapses. Indeed, selective photostimulation of astrocytes with channelrhodopsin-2 in primary visual cortex produced changes in the gain of neuronal tuning similar to those observed in our visual experiments [66]. In the peripheral nervous system, TRPA1 activation at both presynaptic and postsynaptic neurons facilitates excitatory synaptic transmission [67]. In dorsal root ganglia, TRPA1 activation in neurons increases responses to painful and pruritic stimulation. TRPA1 is also expressed in cortical neurons [33] and from previous optovin photoswitching experiments, we know that local activation of TRPA1 in pyramidal neurons can depolarize these cells [13,14]. Taken together, it is therefore possible that the modulations of sensory processing observed here are mediated through both neurons and astrocytes. However, further experiments are needed to identify the relative contribution of neurons and astrocytes to TRPA1 enhancements of sensory processing.

## Methods

### Mice

A total of 22 male wild type and 3 male TRPA1 knockout (KO) C57Bl/6J mice were used in this experiment. All methods were performed in accordance with the protocol approved by the Animal Experimentation and Ethics Committee of the Australian National University (AEEC 2012/64; 2015/74). Mice were housed in a ventilated and air filtered climate-controlled environment with a 12-hour light–dark cycle. Animals had access to food and water ad libitum.

### Surgery for electrophysiological studies

After induction of anesthesia with urethane/chlorprothixene (0.8 g/kg and 5 mg/kg, respectively), mice were placed on a heating blanket (set at ~ 37°C) and the head was fixed in a custom-made apparatus. The scalp was opened through a ~5 mm midline incision and the fascia was removed. A ~2-mm craniotomy was made above the left barrel cortex centered at 1.8 mm posterior and 3.5 mm lateral to Bregma.

### Pipette pair for juxtacellular recording/labelling and pharmacological manipulation

Borosilicate glass patch pipettes were pulled using a micropipette puller (P-97, Sutter Instruments, Novato, CA, USA). For recoding, we pulled the pipettes to reach an impedance of 6–10 MΩ. Then, using a custom-built setup and under microscope guidance, we attached the recording pipette to the infusion pipette (25-50 µm in diameter) with a tip-to-tip distance of ~50 µm (Fig. 1C).

The recording pipette was filled with rat ringer’s solution containing 2% neurobiotin. The infusion pipette was attached to a syringe pump (CMA402, Harvard Apparatus, Holliston, MA, USA) to continuously apply aCSF, AITC or HC at the flow rate of 7 µl/min. The pipette pair was positioned above the craniotomy, and lowered rapidly using a micromanipulator (MPC-200, Sutter Instruments, Novato, CA, USA). After passing through the dura matter, the pressure inside the recording pipette was dropped from ~300 mm Hg to 15–20 mmHg and the pipette pair was advanced at a speed of 2 µm/s while searching for neurons. The resistance was constantly monitored using the current clamp mode of a BVC-700A amplifier (Dagan Corporation, Minneapolis, MN) and through application of 1-nA current pulses with a duration of 200 ms at a frequency of 2.5 Hz. Upon observing fluctuations indicating proximity to a cell and a >4 fold increase in pipette resistance, the pressure was removed and juxtacellular (loose cell-attached) recording was performed. At the end of the recording session, the neuron was loaded with neurobiotin by application of 1–5 nA current pulses of 200-ms duration at a frequency of 2.5 Hz [68].

### Vibrissal stimulation

A Matlab (MathWorks, Inc., Natick, MA) script presented the stimuli and acquired the neuronal data through the analogue input and output of a data acquisition card (National Instruments, Austin, TX) at a sampling rate of 64 kHz. The whisker stimulation protocol was composed of discrete trials of a single deflection at intensities of 0, 25, 50, 100, 150 and 200 µm (250 µm in Ca^2+^ imaging), randomly applied to the contralateral whisker pad. Each deflection was a brief (20 ms) biphasic vertical movement generated by a piezoelectric actuator. The trials were presented in a pseudorandom order at inter-trial intervals of 1 s. Each stimulus was repeated for a minimum of 30 trials. A total of 37 neurons were recorded.

### Visual Stimulation

Drifting grating stimuli were randomly presented at one of 16 different directions. The stimuli were displayed on an 8” monitor (Lilliput 869GL-80NP/C, 1280 x 720 pixel; 60 Hz, Lilliput Ltd., UK) attached to a custom-made funnel and positioned 16 cm from the contralateral (left) eye (Fig. 5A). To apply the stimulus perpendicular to the optic axis, the monitor was rotated 30° relative to the dorsoventral axis of the mouse and tilted 70° off the horizon. The custom-made funnel was used to remove any potentially confounding effect of scattered light from the monitor on the recording camera (Fig. 5A). Visual stimuli were full-screen (58.1° x 32.7°) sinusoidal gratings (0.04 c/°, 80% contrast) drifting at 2 Hz. These values were chosen to be close to the peak visual sensitivity for the largest proportion of mouse V1 neurons [24,69,70]. Parameters of the visual stimuli were selected from the Allen Brain Observatory [69] to measure direction selectivity and were generated in Matlab using the Psychophysics Toolbox [71,72] with a simple power-law gamma correction applied to produce a mid-gray luminance of 24.3 cd/m^2^. Directions from 0° to 337.5° (in 22.5° steps) were presented for 500 ms with a 500 ms inter-stimulus interval. Each direction was presented 35 times in a pseudo-random sequence, along with 35 trials with a blank presentation. A total of 211 neurons were recorded.

### Expression of Ca^2+^ indicator (GCaMP6f)

Mice were briefly anesthetized with isoflurane (~2% by volume in O_2_) in a chamber and moved to a thermal blanket (37°C, Physitemp Instruments) before the head was secured in a stereotaxic frame (Stoelting, IL). Thereafter, the anaesthetic gas (isoflurane, ~2% by volume in O_2_) was passively applied through the nose mask at a flow rate of 0.6-0.8 L/min. The level of anaesthesia was monitored by the respiratory rate, and hind paw and corneal reflexes. The eyes were covered with a thin layer of Viscotears liquid gel (Alcon, UK). The scalp was opened with ~5 mm rostrocaudal incision at the midline using scissor and the periosteum was gently removed. A circular craniotomy was made over the right barrel cortex (3mm diameter; centered 3mm lateral and 1.8mm posterior to Bregma) or the right visual cortex (3mm diameter; centered 2mm lateral and 4.5mm posterior to Bregma) with the dura left intact. A glass pipette (15-25µm diameter at tip) containing GCaMP6f (AAV1.Syn.GCaMP6f.WPRE.SV40, Penn Vector Core, The University of Pennsylvania, USA) was inserted into the cortex at a depth of 230-250 µm below the dura using a micromanipulator (MPC-200, Sutter Instruments, Novato, CA, USA). GCaMP6f was injected at 4-6 sites (at 32nL per site; rate 92 nLs^−1^) using a glass pipette. Injections were controlled using a Nanoject II injector (Drumont scientific, PA). After virus injection, the craniotomy was covered with a 3mm diameter cover-glass (0.1 mm thickness, Warner Instruments, CT). This was glued to the bone surrounding the craniotomy. Custom made head posts were fixed to the skull above Lambda (for somatosensory experiments) or Bregma (for visual experiments) using a thin layer of cyanoacrylate adhesive and dental acrylic. A small well was built surrounding the craniotomy window using dental acrylic to accommodate distilled water required for the immersion lens of the 2-photon microscope.

### Ca^2+^ imaging with 2-photon excitation microscopy

Approximately 3 weeks after injection of GCaMP6f in layer 2/3 cortex, the mouse was anesthetised by intraperitoneal administration of urethane/chlorprothixene (0.8 g/kg and 5 mg/kg body weight, respectively). The mouse was placed on the heating blanket (set at ~ 37°C) and the head was fixed in a custom-built apparatus (Fig. 4A). The 3 mm coverslip was gently removed and a half coverslip was fixed on the medial part of the craniotomy window using dental cement. This allowed us to insert the infusion pipette for pharmacological manipulations during imaging. Ca^2+^ imaging was performed using a two-photon microscope (Thorlabs Inc., Newton, NJ, USA) controlled by ThorImage OCT software. The cortex was illuminated with a Ti:Sapphire fs-pulsed laser (Chameleon, Coherent Inc., Santa Clara, CA, USA) tuned at 920 nm. The laser was focused onto L2/3 cortex through a 16x water-immersion objective lens (0.8NA, Nikon), and Ca^2+^ transients were obtained from neuronal populations at a resolution of 512 × 512 pixels (sampling rate, ~ 30 Hz). To investigate the effect of TRPA1 modulation on neuronal response to whisker or visual stimuli, the recorded neurons were continuously perfused with aCSF, AITC and HC at a speed of 7 µl/min.

For the somatosensory experiments, fluorescence activity was measured after importing images into ImageJ software (National Institutes of Health, USA) and by manual selection of regions of interest (ROIs) around neuronal cell bodies. For the visual experiments, Suite2P[73] was used for motion-correction, ROI-detection and semi-automated ROI selection. We also applied high-(0.1 Hz) and low-pass (29 Hz) filtering to remove low-drifts in the data and high-frequency noise, respectively. For both sensory modalities, the mean background neuropil was subtracted from each neuron’s calcium trace. The change in fluorescence (*ΔF/F0*) was quantified by calculating *F0* either as the mean fluorescence over the whole session (for somatosensory experiments) or 1000 ms window prior to stimulus onset (for visual experiments).

### Immunohistochemistry

At the completion of the experiment, the mouse was euthanised by ip injection of lethabarb 150 mg/kg. Immediately after breathing stopped, the abdomen and chest were opened medially, and the heart was perfused with chilled normal saline. After all blood was washed out of the body; the heart was perfused with 4% paraformaldehyde in phosphate buffered saline (PBS) and the brain was harvested. The brain was fixed in 4% paraformaldehyde in PBS at 4°C overnight. After fixation, the brain was rehydrated gradually by stepwise incubation in 10 to 30% sucrose in PBS (w/v). After rehydration, the brain was sectioned with a Leica CM1580 cryostat (120 µm thickness) and penetrated using PBS containing 1% Triton-X (v/v) for 4–5 h under continuous shaking at room temperature. The slides were then washed three times with PBS. To visualize the neurobiotin loaded neurons, the slices were incubated with streptavidin Alexa Fluor® 488 conjugate (Thermo Fisher Scientific, Waltham, MA, USA) overnight on a shaker at 4°C.

### Quantifying directional selectivity

A permutation procedure was used to determine whether a neuron responded selectively for one direction in the stimulus sequence compared with the response expected by chance. To do this, the difference between post- (500 to 1000 ms) and pre-stimulus (−1000 to 0 ms) was calculated for each trial to determine the stimulus-evoked response. We averaged the evoked responses across all presentations of one direction and compared these with a permuted null distribution (N = 10,000) created by finding the mean response with shuffled labels. The neuron was classified as selective to a given direction if the probability (*p* < 0.0001) of the evoked response from the non-shuffled data was outside (larger or smaller) the values from the shuffled data in each tail of the distribution.

To quantify direction selectivity of the V1 neurons, we fitted the mean stimulus-evoked response (0 to 1000 ms) with a double Gaussian function [25] (Eqn. 1) using non-linear least-squared regression.

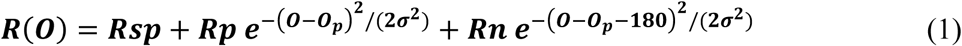

where *O* is the orientation, *Rsp* is the baseline response, *Rp* is the height of the Gaussian at the preferred orientation, *Rn* is the height at the null orientation (preferred + 180°), *O*_*p*_ is the preferred orientation, and σ is the tuning width.

### Linear Discriminant Analysis

We used a multivariate pattern analysis approach to determine the amount of orientation-selective information contained in the population activity. We used the *classify* function in Matlab 2018b with the ‘linear’ option to implement a Linear Discriminant Analysis method. Initially, we applied this at each time point after stimulus presentation to determine the time course of the decoding (Fig. 6B), whereas in later analyses we applied the procedure to time averaged data (500 to 1000 ms, Fig. 6CD) to determine how these effects scale with different numbers of neurons. A 20-fold cross-validation procedure was used to select test and training trials, with each fold having an equal number of trials from each direction. For the initial analysis (Fig. 6B), we used pools of 15 simultaneously-imaged neurons from the same session to account for the noise correlation. For each TRPA1 modulation condition and for each time point (−500 to 2000 ms after stimulus presentation), the classifier was given the presented directions in the training set and the activity of the neurons for those trials, and produced a prediction for the direction presented for the test activity. The trial was classified as correct if the classifier produced the same orientation as was presented in the test set. The cross-validation was repeated 20 times until all trials had been used as test data and for all time points. A new random pool of 15 neurons was then selected and the procedure was repeated for 100 permutations of neurons for each session. This procedure was adapted from previous work on visually-selective neurons [74].

To determine how the classification effects scaled with different population sizes (Fig. 6C) we pooled all recorded neurons (N = 211) from all three sessions and used the mean response from 500 to 1000 ms post-stimulus (i.e., the period over which decoding was maximal in the previous analysis), and used a similar method as described above. The decoding procedure was separately applied to populations from 5 to 205 neurons. For each population size, a random subset of neurons was selected in permutation and was repeated 40 times. To determine whether TRPA1 modulation mainly affected direction or orientation selectivity at a population level, the same procedure was used as in Fig. 6C but with 170 neurons selected for 40 permutations. Now, however, the classifier was trained and tested with either the presented direction or the presented orientation. All other aspects of the analysis were identical to those shown in Fig. 6C.

A standard sign-flipping permutation (N = 10,000) test was used for determining whether decoding accuracy was consistent over time or number of neurons (Fig. 6B). This method makes no assumptions about the underlying shape of the null distribution. This was done by randomly flipping the sign of the data for the decoding permutations with equal probability with a cluster-forming threshold of *p* < 0.05 [75–77] and a cluster significance *p* < 0.05.

## Acknowledgements

We thank Stuart Brierley and Grigori Rychkov for valuable discussions and for providing the TRPA1-KO mice. The experiments were supported by an Australian Research Council (ARC) Discovery Project (DP170100908), an NHMRC project grant (1124411), and the ARC Centre of Excellence for Integrative Brain Function (ARC Centre Grant CE140100007). EA was supported by an ARC Future Fellowship, and JBM was supported by an ARC Australian Laureate Fellowship (FL110100103). The NVIDIA corporation donated a TITAN V GPU to MFT.

